# Gene therapy with doxycycline-controlled expression of human Kv1.1 reduces neuronal excitability and increases sociability of *Scn2a*-deficient mice

**DOI:** 10.64898/2026.09.22.753612

**Authors:** Brody A. Deming, Jingliang Zhang, Iuliia Vitko, Ronald P. Gaykema, Manasi Halurkar, Zaiyang Zhang, Akila D. Abeyaratna, Rineet Ranga, Purba Mandal, Kathryn Lund, Keeley Farmer, Deepti Ramasamy, Edward Perez-Reyes, Yang Yang

**Author notes:** these authors contributed equally to this work (Brody A. Deming, Jingliang Zhang). **To whom correspondence should be addressed:** Yang Yang, PhD; Department of Medicinal Chemistry and Molecular Pharmacology, College of Pharmacy, Purdue University, 207 S Martin Jischke Dr, West Lafayette, IN 47907.

## Abstract

Genetic loss-of-function (LoF) variants in *SCN2A*, a gene encoding the voltage-gated sodium channel Nav1.2, have been identified as one of the foremost monogenic causes of autism spectrum disorder (ASD). ASD encompasses a broad spectrum of behavioral phenotypes, with impaired sociability as a core characteristic. We have established *Scn2a*-deficient mice (*Scn2a^gt/gt^*) to model *Scn2a*-related ASD and found that this model recapitulates social impairment, exhibiting severe social deficits. *Scn2a^gt/gt^* mice exhibit neuronal hyperexcitability and a marked global reduction in potassium channel expression, which plays a crucial role in maintaining the resting membrane potential and repolarizing neurons after an action potential. Among these downregulated potassium channels, potassium voltage-gated channel subfamily A member 1 (Kv1.1) was one of the most affected. To explore whether Kv1.1 could be a potential therapeutic target in *SCN2A*-related ASD, we evaluated the *in vivo* efficacy of a genetic construct driven by the CaMKIIα promoter that allows for exogenous expression of human Kv1.1 (hKv1.1) in principal neurons. In *Scn2a^gt/gt^* mice, we found that hKv1.1 expression normalizes neuronal hyperexcitability. Importantly, doxycycline-induced hKv1.1 expression enhances sociability in *Scn2a*-deficient mice without influencing social behavior in wild-type mice, and this effect was reversed upon doxycycline withdrawal. Overall, we demonstrate the successful use of an inducible AAV-mediated gene delivery system to supplement hKv1.1 expression to mitigate neuronal hyperexcitability and ameliorate social impairments in a mouse model of *SCN2A*-related ASD. These findings highlight the contribution of Kv1.1 to *SCN2A*-associated pathophysiology and its potential as a therapeutic target for severe social deficits.

## INTRODUCTION

Autism Spectrum Disorder (ASD) is one of the most prevalent neurodevelopmental disorders, with CDC estimates indicating that ASD affects approximately 1 in 31 children in the United States [1]. ASD encompasses a spectrum of behavioral abnormalities, with social deficits being a core symptom. ASD is a complex disorder that has been linked to both genetic and environmental factors [2]. While it has generally been thought that ASD is a polygenic disorder [3], mutations in single genes have been linked to the development of ASD [4–6]. It has recently been identified that genetic variants in the gene *SCN2A,* which encodes the voltage-gated sodium ion channel alpha subunit II (Nav1.2), are one of the leading monogenic causes of ASD [7]. Genetic variants in *SCN2A* generally fall into two categories: gain-of-function (GoF) and loss-of-function (LoF), with GoF mutations typically manifesting in epilepsy and LoF mutations leading to the development of ASD. While sodium channel blockers can be used to treat epilepsy related to Nav1.2 GoF, treatments for *SCN2A* LoF-related ASD are lacking.

To study *SCN2A* LoF-related ASD and test potential interventions, establishing an appropriate animal model is essential. Recapitulating behavioral phenotypes of ASD in animal models has been challenging [8]. Specifically, *Scn2a^+/-^* mice expressing 50% of wild-type (WT) Nav1.2 display mild behavioral abnormalities and show no significant social impairments, while *Scn2a^-/-^* mice fail to survive postnatally [9–13]. To capture key behavioral features of severe *SCN2A* LoF, we previously developed a gene-trap mouse model by inserting a trapping cassette into the *Scn2a* locus (*Scn2a^gt/gt^*), which reduces *Scn2a* mRNA and protein expression to less than half of WT levels. The resulting mice are viable and exhibit core phenotypes of ASD [14–17]. We also detected paradoxical neuronal hyperexcitability in the *Scn2a^gt/gt^* mice [18], together with downregulation of multiple potassium channels, including *Kcna1*, suggesting a possible compensatory mechanism [18].

Interestingly, Kv1.1 is a promising target for the development of gene therapies due to its crucial role in regulating neuronal excitability and repolarization following an action potential (AP) [19]. LoF mutations in *KCNA1* cause severe epilepsy and developmental disability, motivating the exploration of gene therapy approaches to ameliorate these phenotypes [20, 21]. Importantly, studies have shown that gene therapy to deliver exogenous *KCNA1* is an effective approach to reduce neuronal hyperexcitability and alleviate epileptic seizures [20, 22, 23]. While gene therapy approaches have shown great promise, enhancing their precision and tunability remains an area of active development. Adjustable gene expression can be achieved by using a tetracycline-inducible transactivator system, allowing gene dosage to be modulated by varying doxycycline administration [24]. The Tet-On system would allow for a more personalized gene therapy approach and would be beneficial in disorders with heterogeneous patient phenotypes, such as *SCN2A*-related ASD.

In this study, we evaluated whether enhancing Kv1.1 expression is sufficient to alleviate electrophysiological abnormalities and rescue social deficits in *Scn2a^gt/gt^* mice. Given the critical involvement of the striatum and medial prefrontal cortex (mPFC) in ASD [11, 25–27], we examined how exogenous expression of human Kv1.1 (hKv1.1) alters the intrinsic electrophysiological properties of striatal medium spiny neurons (MSNs) and Layer V pyramidal neurons in the mPFC. We confirmed that *Scn2a^gt/gt^* mice exhibited abnormal intrinsic electrophysiological properties consistent with our previous findings [18], and that hKv1.1 expression restored their intrinsic electrophysiological properties. Importantly, we found that exogenous expression of hKv1.1 alleviated social deficits in *Scn2a^gt/gt^* mice. The doxycycline-inducible Tet-On system in the AAV-hKv1.1 expression cassette allowed us to assess the reversibility of the behavioral effects; after doxycycline administration was stopped, social behavior returned to baseline. This study sheds light on the role Kv1.1 plays in *SCN2A*-related ASD and demonstrates its potential as a therapeutic target.

## RESULTS

### Doxycycline administration controls functional hKv1.1 channel expression *in vitro* and *in vivo*

Recent developments in gene therapy have highlighted the potential therapeutic benefit of targeting Kv1.1 in epilepsy [20, 28]. Gene therapies cannot typically be tuned after delivery, whereas small molecules allow physicians to adjust dosages and provide a more personalized treatment approach. With this in mind, we have developed a genetic construct that utilizes a tetracycline-controlled Tet-On transcriptional activation system to express hKv1.1 (**Figure 1A**). This system allows for constitutive expression of a reverse tetracycline transactivator (rtTA) (Tet-On 3G). This generation of rtTA has been optimized for doxycycline-induced expression and diminished background activity [24]. To validate that this construct leads to functional hKv1.1 expression, plasmids were transfected into ND7/23 rodent neuronal cells, and whole-cell patch-clamp recordings were performed to record potassium currents in the presence or absence of doxycycline (**Figure 1B**). We found that transfected ND7/23 cells produced robust potassium currents upon doxycycline treatment, whereas vehicle-treated cells did not (**Figure 1C**). To further validate this genetic construct *in vivo*, we packaged the construct into an adeno-associated virus (AAV) and injected it into the striatum and medial prefrontal cortex (mPFC) of WT mice. qPCR and Western blot analyses were then performed after 2 weeks of doxycycline or control chow administration.

**Figure 1:**
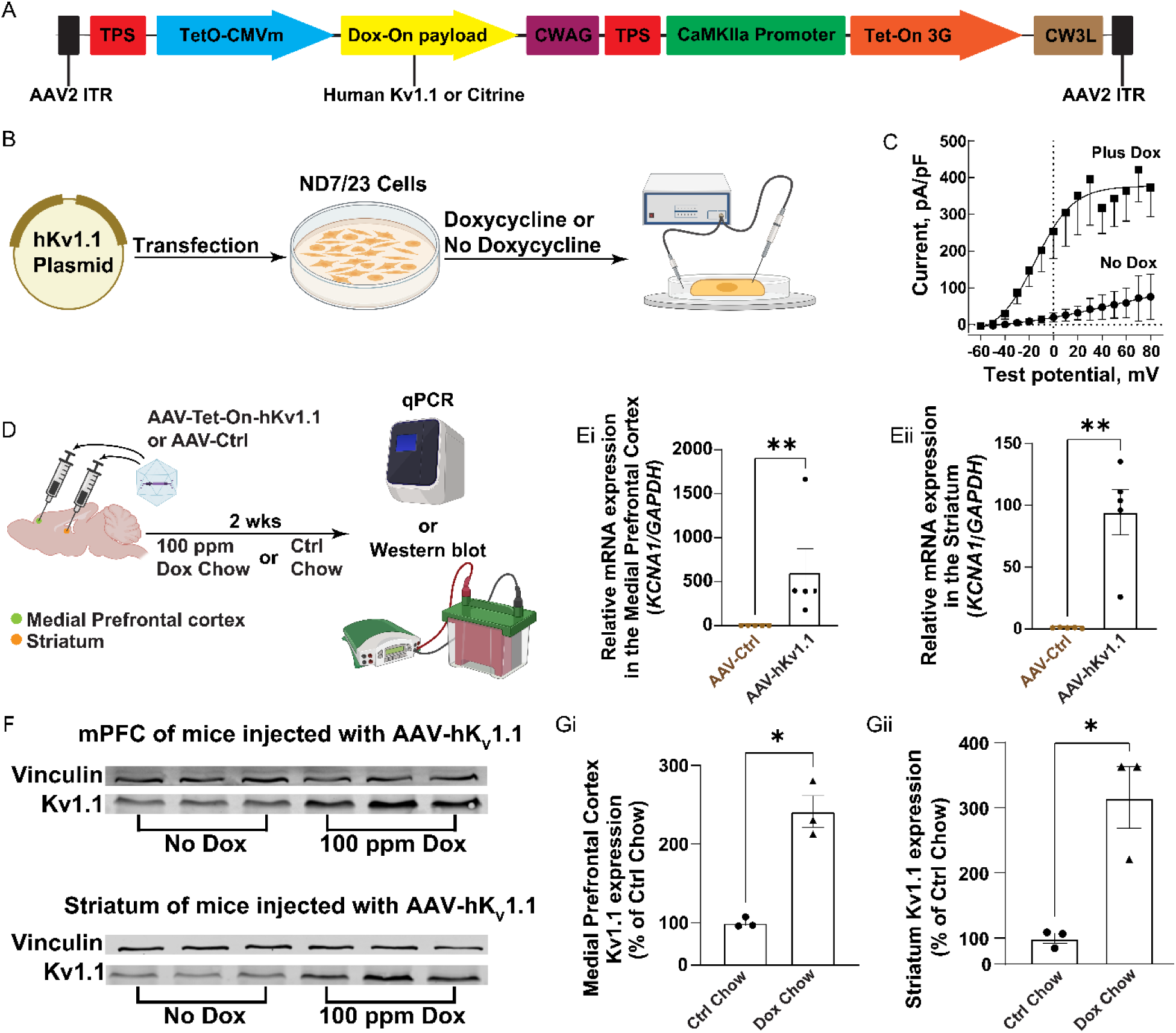
Doxycycline administration leads to functional hKv1.1 channels *in vitro* and hKv1.1 expression *in vivo*. (A) Plasmid backbone of Tet-On system: The black regions are Internal Terminal Repeats (ITR), the red regions are Transcriptional Pause Sites (TPS), the blue region is the tet-operator (TetO-CMVm), the yellow region is the Doxycycline payload (Dox-On payload), the maroon region is the (CWAG) sequence, which reduces leaky expression, the green region is the CaMKII promoter (CaMKII Promoter), the orange region is the third-generation reverse tetracycline transactivator (Tet-On 3G), the brown region is the minimal woodchuck post-regulatory element used for efficient expression. (B) Schematic of experimental design for whole-cell patch-clamp recordings performed on ND7/23 cells transfected with the hKv1.1 plasmid. (C) Current-voltage relationship normalized to cell capacitance in ND7/23 cells transfected with the hKv1.1 plasmid and treated with either doxycycline (n=9) or no doxycycline (n=5). The currents in the No Dox condition are due to background K+ currents. (D) Schematic of experimental design for viral injections. (Ei) Quantification of relative mRNA expression of *KCNA1* in the medial prefrontal cortex of mice injected with AAV-Ctrl and AAV-hKv1.1 and fed 100 ppm dox chow. n= WT-Ctrl: 5, WT-hKv1.1: 5. Mann-Whitney test: **, p<0.01. (Eii) Quantification of relative mRNA expression of *KCNA1* in the striatum of mice injected with AAV-Ctrl and AAV-hKv1.1 and fed 100 ppm dox chow. n= WT-Ctrl: 5, WT-hKv1.1: 5. Mann-Whitney test: **, p<0.01. (F) Representative Western blot images showing the medial prefrontal cortex and striatum of mice injected with AAV-hKv1.1 and fed 100 ppm dox chow. (Gi) Quantification of Kv1.1 expression in the medial prefrontal cortex of mice injected with AAV-hKv1.1 and fed 100 ppm dox chow. n= WT-hKv1.1+Ctrl-chow: 3, WT-hKv1.1+Dox-chow: 3. Welch’s t-test: *, p<0.05. (Gii) Quantification of Kv1.1 expression in the striatum of mice injected with AAV-hKv1.1 and fed 100 ppm dox chow. n= WT-hKv1.1+Ctrl-chow: 3, WT-hKv1.1+Dox-chow: 3. Welch’s t-test: *, p<0.05.

qPCR analysis confirmed that hKv1.1 mRNA was detected only in mice injected with AAV-hKv1.1 and not in mice injected with AAV-Ctrl, despite both groups being fed doxycycline chow (**Figure 1Ei and Eii**). Upon western blot analysis, we found that mice injected with AAV-hKv1.1 and fed doxycycline had a significant increase in Kv1.1 protein expression, while those maintained on control chow showed no change (**Figure 1Gi and Gii**). Our *in vitro* and *in vivo* findings confirmed that the hKv1.1 Tet-On system drives functional hKv1.1 channel expression only in the presence of doxycycline.

### Exogenous expression of hKv1.1 in the striatum decreases hyperexcitability observed in medium spiny neurons of *Scn2a*-deficient mice

To test whether exogenous expression of hKv1.1 can rescue the electrophysiological abnormalities observed in *SCN2A*-deficient mice, we injected AAV-hKv1.1 or AAV-Ctrl into the striatum of *Scn2a^gt/gt^* and WT mice, a brain region that has repeatedly been shown to be affected in ASD [25, 26, 29]. After doxycycline-induced hKv1.1 expression, we conducted whole-cell patch-clamp recordings on putative medium spiny neurons (MSNs), which constitute ~95% of the striatal neuron population and exhibit intrinsic hyperexcitability in *Scn2a^gt/gt^* mice [18]. We observed that *Scn2a^gt/gt^-*Ctrl mice had significantly increased AP firing, while *Scn2a^gt/gt^*-hKv1.1 mice exhibited decreased AP firing during a step current injection of 50-400 pA, similar to that of WT-Ctrl mice (**Figure 2C-D**). We then evaluated the effects of exogenous hKv1.1 expression on resting membrane potential (RMP), input resistance, rheobase, and action potential amplitude. We also measured δ-after-hyperpolarization (δ-AHP) to account for potential differences in RMP. We found that exogenous expression of hKv1.1 in *Scn2a^gt/gt^* led to a decrease in RMP (*Scn2a^gt/gt^*-hKv1.1: −87.45 ± 0.3721 mV and *Scn2a^gt/gt^*-Ctrl: −85.35 ± 0.4750 mV), a decrease in input resistance (*Scn2a^gt/gt^*-hKv1.1: 114.9 ± 8.156 MΩ and *Scn2a^gt/gt^*-Ctrl: 146.2 ± 8.109 MΩ), and an increase in rheobase (*Scn2a^gt/gt^*-hKv1.1: 574.7 ± 23.93 pA and *Scn2a^gt/gt^*-Ctrl: 465.3 ± 20.00 pA) compared to *Scn2a^gt/gt^*-Ctrl mice. We found that the electrophysiological changes observed in *Scn2a^gt/gt^*-hKv1.1 mice were similar to the electrophysiological properties of MSNs in WT-Ctrl mice (WT-Ctrl mice: RMP: −87.70 ± 0.5406 mV, input resistance: 90.16 ± 7.391 MΩ, rheobase: 580.7 ± 22.69 pA) (**Figure 2F-J**). While exogenous expression of hKv1.1 led to changes in many electrophysiological parameters of MSNs, the amplitude of the APs (WT-Ctrl: 111.20 ± 1.122 mV, *Scn2a^gt/gt^-*Ctrl: 102.80 ± 0.7911 mV, and *Scn2a^gt/gt^*-hKv1.1: 104.80 ± 1.248 mV) and δ-AHP (WT-Ctrl: 33.28 ± 0.7554 mV, *Scn2a^gt/gt^*-Ctrl: 33.95 ± 0.5200 mV, and *Scn2a^gt/gt^*-hKv1.1: 35.39 ± 0.8449 mV) of MSNs remained unchanged in *Scn2a^gt/gt^*-hKv1.1 mice (**Figure 2K-L**).

**Figure 2:**
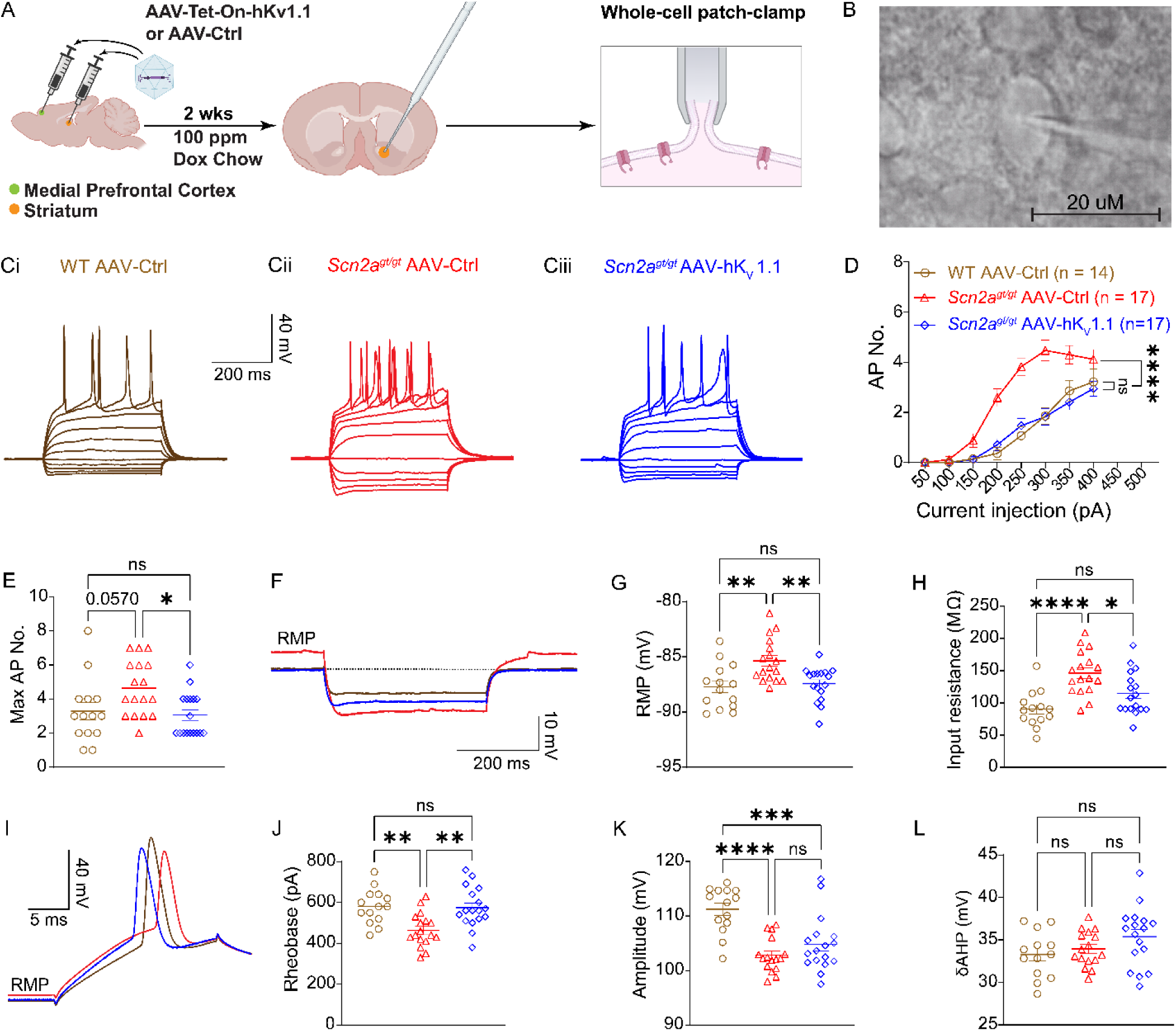
Exogenous expression of hKv1.1 in the striatum decreases hyperexcitability observed in medium spiny neurons of *Scn2a*-deficient mice. (A) Schematic of experimental design. (B) Representative image of a typical MSN patched during the experiment. (Ci-Ciii) Representative current-clamp recordings obtained at resting membrane potential from MSNs in the nucleus accumbens of WT mice injected with AAV-Ctrl (brown) and *Scn2a*-deficient mice injected with AAV-Ctrl (red) or AAV-hKv1.1 (blue). (D) Number of action potentials of MSNs from WT mice injected with AAV-Ctrl (brown) and *Scn2a*-deficient mice injected with AAV-Ctrl (red) or AAV-hKv1.1 (blue) generated in response to depolarizing current pulses (50-400 pA). Two-way ANOVA with Tukey’s multiple comparisons: ns= not significant, p>0.05; ****, p<0.0001. (E) Maximum number of action potentials of MSNs from WT mice injected with AAV-Ctrl (brown) and *Scn2a*-deficient mice injected with AAV-Ctrl (red) or AAV-hKv1.1 (blue) generated in response to depolarizing current pulses (50-400 pA). Kruskal-Wallis with Dunn’s multiple comparisons: ns= not significant, p>0.05; *, p<0.05. (F) Representative traces of MSNs from WT mice injected with AAV-Ctrl (brown) and *Scn2a*-deficient mice injected with AAV-Ctrl (red) or AAV-hKv1.1 (blue) generated in response to a −100 pA current injection. (G) Resting membrane potential of MSNs from WT mice injected with AAV-Ctrl (brown) and *Scn2a*-deficient mice injected with AAV-Ctrl (red) or AAV-hKv1.1 (blue). One-way ANOVA with Tukey’s multiple comparisons: ns= not significant, p>0.05; **, p<0.01. (H) Input resistance of MSNs from WT mice injected with AAV-Ctrl (brown) and *Scn2a*-deficient mice injected with AAV-Ctrl (red) or AAV-hKv1.1 (blue). One-way ANOVA with Tukey’s multiple comparisons: ns= not significant, p>0.05; *, p<0.05; ****, p<0.0001. (I) Representative action potential from MSNs of WT mice injected with AAV-Ctrl (brown) and *Scn2a*-deficient mice injected with AAV-Ctrl (red) or AAV-hKv1.1 (blue). (J) Rheobase of MSNs from WT mice injected with AAV-Ctrl (brown) and *Scn2a*-deficient mice injected with AAV-Ctrl (red) or AAV-hKv1.1 (blue). One-way ANOVA with Tukey’s multiple comparisons: ns= not significant, p>0.05; **, p<0.01. (K) Action potential amplitude of MSNs from WT mice injected with AAV-Ctrl (brown) and *Scn2a*-deficient mice injected with AAV-Ctrl (red) or AAV-hKv1.1 (blue). One-way ANOVA with Tukey’s multiple comparisons: ns= not significant, p>0.05; ****, p<0.0001. (L) δ-fast-after-hyperpolarization (δ-AHP) of action potential spikes of MSNs from WT mice injected with AAV-Ctrl (brown) and *Scn2a*-deficient mice injected with AAV-Ctrl (red) or AAV-hKv1.1 (blue). One-way ANOVA with Tukey’s multiple comparisons: ns= not significant.

To determine how RMP influenced these findings, we fixed the neurons at −80 mV and performed another set of whole-cell patch-clamp recordings. We found that MSNs of *Scn2a^gt/gt^*-Ctrl mice still exhibited hyperexcitability, indicated by an increased number of APs during a current injection of 50-400 pA (**Supplementary Figure 1A-B**). *Scn2a^gt/gt^*-hKv1.1 mice also showed an increase in rheobase to a level comparable to that of WT-Ctrl mice (WT-Ctrl: 492.1 ± 17.95 pA, *Scn2a^gt/gt^*-Ctrl: 421.2 ± 16.62 pA, and *Scn2a^gt/gt^*-hKv1.1: 501.8 ± 24.48 pA) (**Supplementary Figure 1G**). Together, these results suggest that the intrinsic hyperexcitability observed in *Scn2a^gt/gt^*-Ctrl mice is not solely due to a depolarized RMP. Our data also demonstrate that exogenous expression of hKv1.1 is sufficient to normalize abnormal passive membrane properties in striatal neurons of *Scn2a^gt/gt^* mice.

### Exogenous expression of hKv1.1 in the medial prefrontal cortex decreases hyperexcitability observed in Layer V pyramidal neurons of *Scn2a*-deficient mice

Given that hyperexcitability is not just observed in striatal MSNs of *Scn2a^gt/gt^* mice, we next asked whether hKv1.1 expression could reverse the hyperexcitability previously observed in Layer V pyramidal neurons of the mPFC [11, 18]. To address this question, we injected AAV-hKv1.1 into the mPFC of *Scn2a*^gt/gt^ mice and performed whole-cell patch-clamp recordings to evaluate whether exogenous expression of hKv1.1 could also rescue the intrinsic hyperexcitability of Layer V pyramidal neurons. We found that *Scn2a^gt/gt^*-Ctrl mice exhibited an increase in the number of APs fired during a current injection of 50-400 pA (**Figure 3C-D**). Furthermore, we found that Layer V pyramidal neurons in *Scn2a^gt/gt^*-Ctrl mice had an increase in RMP (*Scn2a^gt/gt^*-Ctrl: −68.97 ± 0.7378 mV) and input resistance (*Scn2a^gt/gt^*-Ctrl: 179.3 ± 16.57 MΩ) (**Figure 3G-H**); Layer V pyramidal neurons in *Scn2a^gt/gt^-*Ctrl mice also had a decrease in rheobase (*Scn2a^gt/gt^*-Ctrl: 308.8 ± 20.59 pA), amplitude (*Scn2a^gt/gt^*-Ctrl: 91.97 ± 1.036 mV), and δ-AHP (*Scn2a^gt/gt^*-Ctrl: 20.73 ± 0.8278 mV) (**Figure 3J-L**). *Scn2a^gt/gt^*-hKv1.1 mice had a significant decrease in the number of APs fired during a current injection of 50-400 pA (**Figure 3C-D**). It was also observed that Layer V pyramidal neurons of *Scn2a^gt/gt^*-hKv1.1 mice had a decreased RMP (*Scn2a^gt/gt^*-hKv1.1: −73.43 ± 0.5393 mV) and input resistance (*Scn2a^gt/gt^*-hKv1.1: 135.0 ± 9.281 MΩ). Moreover, *Scn2a^gt/gt^*-hKv1.1 mice exhibited significant increases in rheobase (*Scn2a^gt/gt^*-hKv1.1: 397.8 ± 19.18 pA), amplitude (*Scn2a^gt/gt^*-hKv1.1: 97.59 ± 0.9380 mV), and δ-AHP (*Scn2a^gt/gt^*-hKv1.1: 25.70 ± 0.5987 mV) (**Figure 3J-L**). The electrophysiological changes observed in *Scn2a^gt/gt^*-hKv1.1 mice were similar to those of WT-Ctrl mice (WT-Ctrl: RMP: −74.88 ± 0.6856 mV, input resistance: 104.1 ± 11.89 MΩ, rheobase: 504.5 ± 20.59 pA, amplitude: 106.2 ± 0.7729 mV, and δ-AHP: 26.60 ± 0.7962 mV). These results demonstrate that exogenous expression of hKv1.1 alters the intrinsic electrophysiological properties of Layer V pyramidal neurons of the PFC in *Scn2a^gt/gt^* mice.

**Figure 3:**
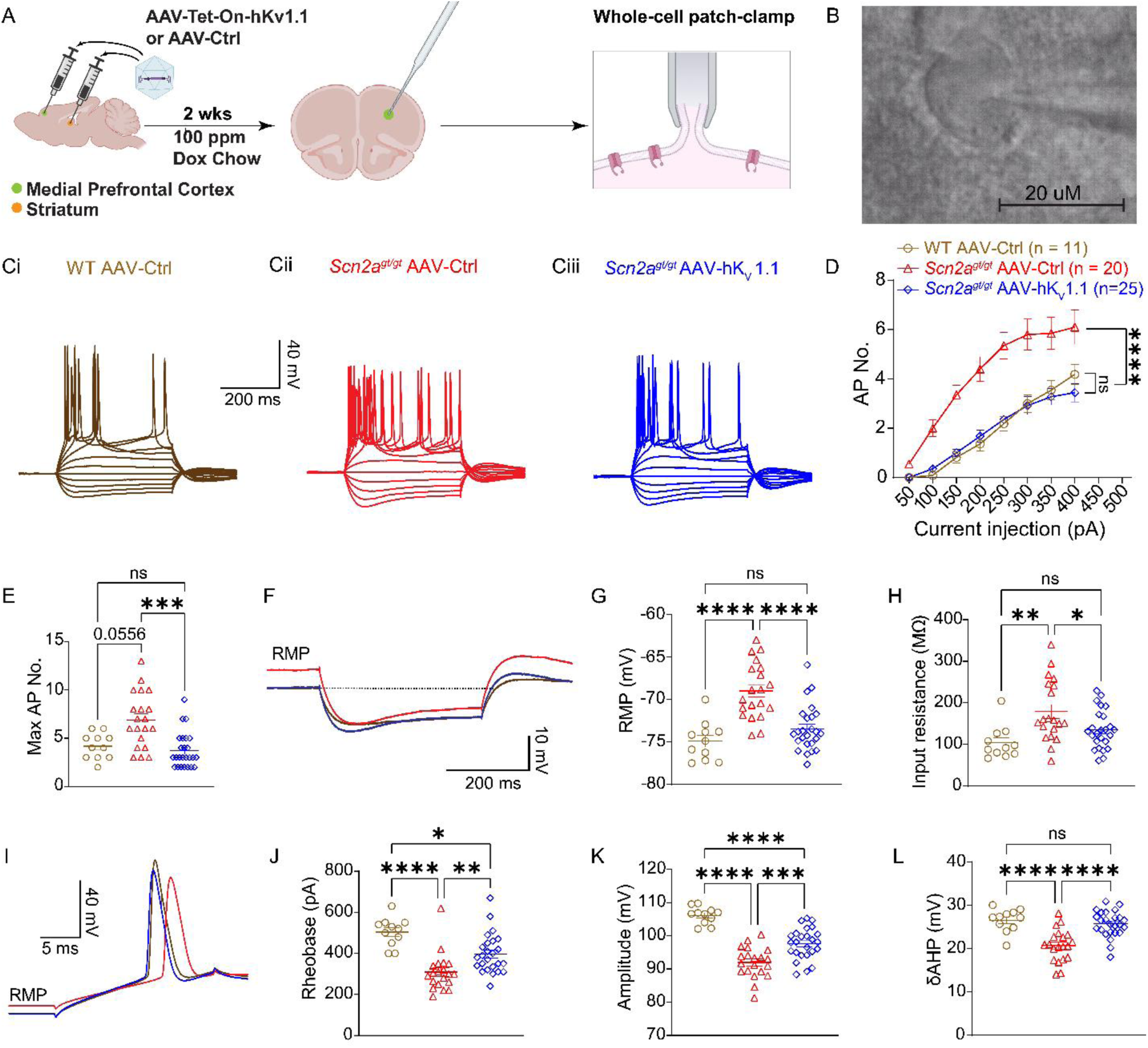
Exogenous expression of hKv1.1 in the medial prefrontal cortex decreases hyperexcitability observed in Layer V pyramidal neurons of *Scn2a*-deficient mice. (A) Schematic of experimental design. (B) Representative image of a typical Layer V pyramidal neuron patched during the experiment. (Ci-Ciii) Representative current-clamp recordings of Layer V pyramidal neurons obtained at resting membrane potential from WT mice injected with AAV-Ctrl (brown) and *Scn2a*-deficient mice injected with AAV-Ctrl (red) or AAV-hKv1.1 (blue). (D) Number of action potentials of Layer V pyramidal neurons from WT mice injected with AAV-Ctrl (brown) and *Scn2a*-deficient mice injected with AAV-Ctrl (red) or AAV-hKv1.1 (blue) generated in response to depolarizing current pulses (50-400 pA). Two-way ANOVA with Tukey’s multiple comparisons: ns= not significant, p>0.05; ***, p<0.001. (E) Maximum number of action potentials of Layer V pyramidal neurons from WT mice injected with AAV-Ctrl (brown) and *Scn2a*-deficient mice injected with AAV-Ctrl (red) or AAV-hKv1.1 (blue) generated in response to depolarizing current pulses (50-400 pA). Kruskal-Wallis with Dunn’s multiple comparisons: ns= not significant, p>0.05; *, p<0.05. (F) Representative traces of Layer V pyramidal neurons from WT mice injected with AAV-Ctrl (brown) and *Scn2a*-deficient mice injected with AAV-Ctrl (red) or AAV-hKv1.1 (blue) generated in response to a −100 pA current injection. (G) Resting membrane potential of Layer V pyramidal neurons from WT mice injected with AAV-Ctrl (brown) and *Scn2a*-deficient mice injected with AAV-Ctrl (red) or AAV-hKv1.1 (blue). One-way ANOVA with Tukey’s multiple comparisons: ns= not significant, p>0.05; ****, p<0.0001. (H) Input resistance of Layer V pyramidal neurons from WT mice injected with AAV-Ctrl (brown) and *Scn2a*-deficient mice injected with AAV-Ctrl (red) or AAV-hKv1.1 (blue). One-way ANOVA with Tukey’s multiple comparisons: ns= not significant, p>0.05; *, p<0.05; **, p<0.01. (I) Representative action potential from Layer V pyramidal neurons of WT mice injected with AAV-Ctrl (brown) and *Scn2a*-deficient mice injected with AAV-Ctrl (red) or AAV-hKv1.1 (blue). (J) Rheobase of Layer V pyramidal neurons from WT mice injected with AAV-Ctrl (brown) and *Scn2a*-deficient mice injected with AAV-Ctrl (red) or AAV-hKv1.1 (blue). One-way ANOVA with Tukey’s multiple comparisons: *, p<0.05; **, p<0.01; ***, p<0.001. (K) Action potential amplitude of Layer V pyramidal neurons from WT mice injected with AAV-Ctrl (brown) and *Scn2a*-deficient mice injected with AAV-Ctrl (red) or AAV-hKv1.1 (blue). One-way ANOVA with Tukey’s multiple comparisons: ***, p<0.001; ****, p<0.0001. (L) δ-fast-after-hyperpolarization (δ-AHP) of action potential spikes of Layer V pyramidal neurons from WT mice injected with AAV-Ctrl (brown) and *Scn2a*-deficient mice injected with AAV-Ctrl (red) or AAV-hKv1.1 (blue). One-way ANOVA with Tukey’s multiple comparisons: ns= not significant; ****, p<0.0001.

To evaluate what role RMP plays in the hyperexcitability of Layer V pyramidal neurons, we fixed these cells at a membrane potential of −70 mV and performed whole-cell patch-clamp recordings. We found that exogenous expression of hKv1.1 could lower the number of APs fired (**Supplementary Figure 2A-B**) while input resistance (WT-Ctrl: 114.1 ± 11.15 MΩ, *Scn2a*^gt/gt^-Ctrl: 166.0 ± 14.89 MΩ, and *Scn2a*^gt/gt^-hKv1.1: 144.7 ± 9.821 MΩ), amplitude (WT-Ctrl: 100.5 ± 0.9466 mV, *Scn2a*^gt/gt^-Ctrl: 92.15 ± 0.8444 mV, and *Scn2a*^gt/gt^-hKv1.1: 93.62 ± 0.6899 mV), AHP (WT-Ctrl: −46.76 ± 0.6062 mV, *Scn2a*^gt/gt^-Ctrl: −47.09 ± 0.4390 mV, and *Scn2a*^gt/gt^-hKv1.1: −46.93 ± 0.4566 mV), and half-width (WT-Ctrl: 2.501 ± 0.07260 ms, *Scn2a*^gt/gt^-Ctrl: 2.589 ± 0.05808 ms, and *Scn2a*^gt/gt^-hKv1.1: 2.684 ± 0.06922 ms), remained unchanged (**Supplementary Figure 2C-J**), confirming that RMP plays an important part in the hyperexcitability observed in Layer V pyramidal neurons of *Scn2a^gt/gt^* mice.

### Local exogenous expression of hKv1.1 in the prefrontal cortex and striatum increases the sociability of *Scn2a*-deficient mice

To evaluate if exogenous expression of hKv1.1 could mitigate behavioral impairments previously observed in *Scn2a^gt/gt^* mice [14, 16], we injected AAV-Ctrl or AAV-hKv1.1 bilaterally into the striatum and mPFC of WT and *Scn2a^gt/gt^* mice. Taking advantage of the Tet-On system, we temporally controlled hKv1.1 expression and assessed sociability in the three-chamber assay under control chow, doxycycline chow, and washout conditions (**Figure 4A**). During the three-chamber assay, we observed that *Scn2a^gt/gt^* mice injected with either AAV-Ctrl or AAV-hKv1.1 spent less time in the chamber with a novel mouse prior to doxycycline administration (*Scn2a*^gt/gt^-Ctrl: 62.06 ± 24.80 s, *Scn2a*^gt/gt^-hKv1.1: 30.03 ± 14.03 s). When doxycycline was administered, we observed that *Scn2a*^gt/gt^-hKv1.1 mice had a significant increase in the time spent in the social chamber with the novel mouse, suggesting that doxycycline-induced expression of hKv1.1 in the mPFC and striatum results in increased social behavior of *Scn2a^gt/gt^* mice (*Scn2a*^gt/gt^-Ctrl: 26.11 ± 14.09 s, *Scn2a*^gt/gt^-hKv1.1: 101.2 ± 25.27 s). Importantly, sociability decreased following doxycycline withdrawal, supporting the conclusion that the increase in sociability observed in *Scn2a^gt/gt^*-hKv1.1 while on Dox chow was dependent on doxycycline-induced hKv1.1 expression (*Scn2a*^gt/gt^-Ctrl: 42.10 ± 19.81 s, *Scn2a*^gt/gt^-hKv1.1: 54.07 ± 17.40 s) (**Figure 4B-G**).

**Figure 4:**
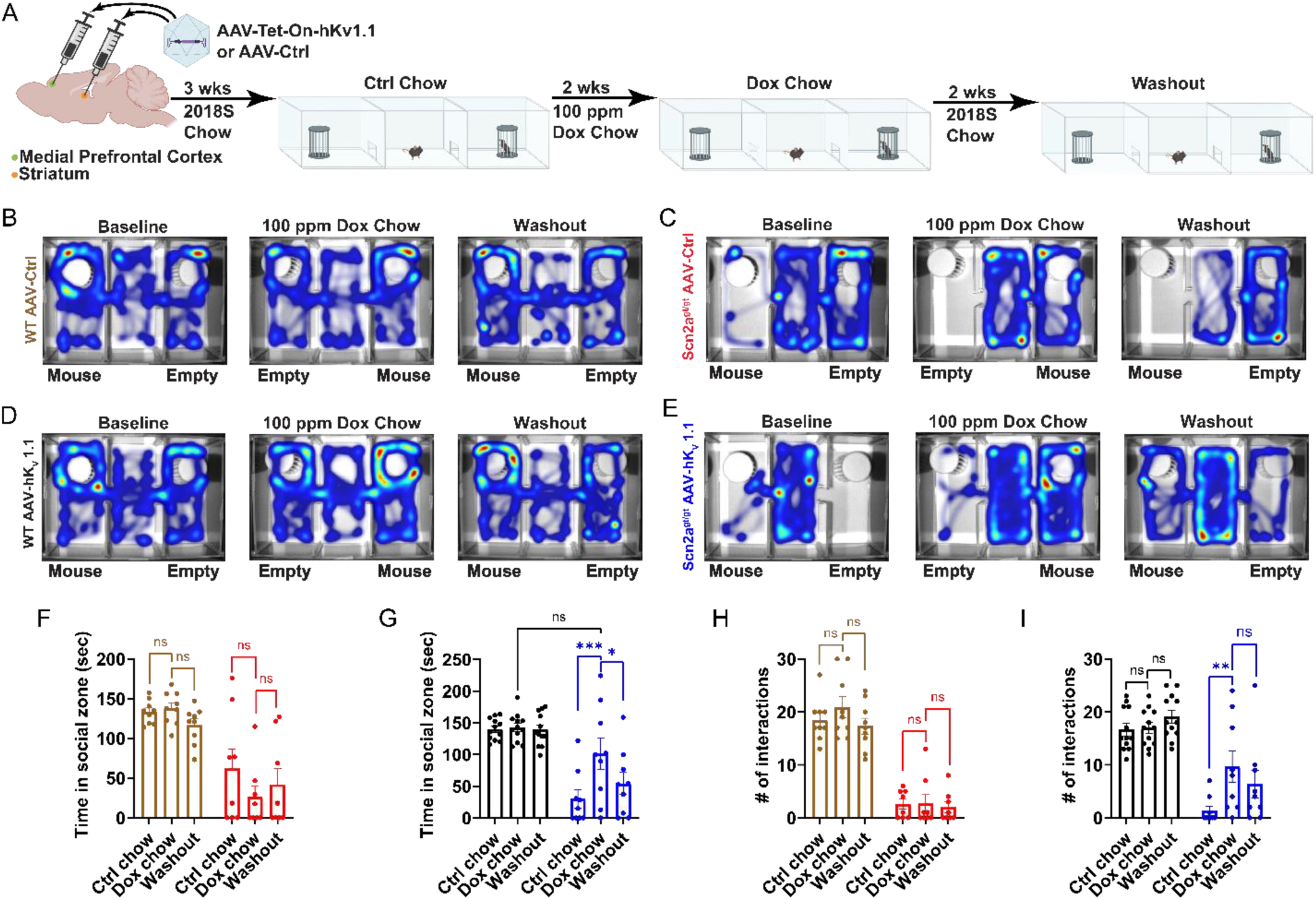
Local exogenous expression of hKv1.1 in the medial prefrontal cortex and striatum increases sociability of *Scn2a*-deficient mice. (A) Schematic of experimental design. (B) Representative heatmaps from WT mice injected with AAV-Ctrl during a 5-min three-chamber assay following control chow (baseline), 100 ppm doxycycline chow (Dox chow), and control chow after doxycycline withdrawal (washout). (C) Representative heatmaps of *Scn2a*-deficient mice injected with AAV-Ctrl during a 5-min three-chamber assay following control chow (baseline), 100 ppm doxycycline chow (Dox chow), and control chow after doxycycline withdrawal (washout). (D) Representative heatmaps of WT mice injected with AAV-hKv1.1 during a 5-min three-chamber assay following control chow (baseline), 100 ppm doxycycline chow (Dox chow), and control chow after doxycycline withdrawal (washout). (E) Representative heatmaps of *Scn2a*-deficient mice injected with AAV-hKv1.1 during a 5-min three-chamber assay following control chow (baseline), 100 ppm doxycycline chow (Dox chow), and control chow after doxycycline withdrawal (washout). (F) Time WT and *Scn2a*-deficient mice injected with AAV-Ctrl spent in the social zone during a 5-minute three-chamber assay. n= WT-Ctrl: 9, *Scn2a*-Ctrl: 8. Two-way repeated-measures ANOVA with Bonferroni multiple comparisons: ns= not significant, p>0.05. (G) Time WT and *Scn2a*-deficient mice injected with AAV-hKv1.1 spent in the social zone during a 5-minute three-chamber assay. n= WT-hKv1.1: 12, *Scn2a*-hKv1.1: 9. Two-way repeated-measures ANOVA with Bonferroni multiple comparisons: ns= not significant, p>0.05; *, p<0.05; ***, p<0.001. (H) Number of interactions WT and *Scn2a*-deficient mice injected with AAV-Ctrl had with a novel mouse during a 5-minute three-chamber assay. n= WT-Ctrl: 9, *Scn2a*-Ctrl: 8. Two-way repeated-measures ANOVA with Bonferroni multiple comparisons: ns= not significant, p>0.05. (I) Number of interactions WT and *Scn2a*-deficient mice injected with AAV-hKv1.1 had with a novel mouse during a 5-minute three-chamber assay. n= WT-hKv1.1: 12, *Scn2a*-hKv1.1: 9. Two-way repeated-measures ANOVA with Bonferroni multiple comparisons: ns= not significant, p>0.05; *, p<0.05; **, p<0.01.

Upon further analysis of the three-chamber videos, we observed that *Scn2a*^gt/gt^-hKv1.1 mice receiving doxycycline were not just spending more time in the social chamber but also interacted more frequently with the novel mouse, which was not observed in *Scn2a*^gt/gt^-Ctrl mice (Ctrl chow: *Scn2a*^gt/gt^-Ctrl: 2.625 ± 0.9246, *Scn2a*^gt/gt^-hKv1.1: 1.333 ± 0.8333, 100 ppm Dox chow: *Scn2a*^gt/gt^-Ctrl: 2.750 ± 1.688, *Scn2a*^gt/gt^-hKv1.1: 9.667 ± 2.967, Washout: *Scn2a*^gt/gt^-Ctrl: 2.000 ± 1.018, *Scn2a*^gt/gt^-hKv1.1: 6.333 ± 2.619) (**Figure 4H-I**). To determine whether the increase in social behavior was associated with an increase in activity level, we performed an open field assay following doxycycline administration. We found no significant difference in the distance traveled (*Scn2a*^gt/gt^-Ctrl: 5284 ± 1085 cm, and *Scn2a*^gt/gt^-hKv1.1: 6213 ± 1326 cm) or velocity (*Scn2a*^gt/gt^-Ctrl: 8.807 ± 1.809 cm/s, and *Scn2a*^gt/gt^-hKv1.1: 10.35 ± 2.210 cm/s) (**Supplementary Figure 3A, D and E**) of *Scn2a*^gt/gt^ mice injected with AAV-Ctrl or AAV-hKv1.1, suggesting that hKv1.1 expression did not significantly alter locomotor activity. Together, this data shows that enhancing Kv1.1 can rescue social deficits observed in *Scn2a^gt/gt^ mice*.

*Scn2a*-deficient mice also present with innate behavioral deficits, such as the lack of nest building and decreased marble burying [14]. However, the interpretation of these behaviors can be complex, as they are not specific to a single behavioral phenotype [30, 31]. Nevertheless, we evaluated whether exogenous expression of hKv1.1 could also rescue these innate behavioral abnormalities in *Scn2a*^gt/gt^ mice. We found that adult *Scn2a*^gt/gt^ mice injected with AAV-Ctrl or AAV-hKv1.1 did not significantly differ in the number of marbles buried or the quality of the nest built. In both groups, 100% of the marbles remained unburied after 30 minutes (*Scn2a*^gt/gt^-Ctrl: 0.000 ± 0.000, and *Scn2a*^gt/gt^-hKv1.1: 0.000 ± 0.0000) (**Supplementary Figure 3F-H**), and most of the nesting material remained untouched after 36 h (*Scn2a*^gt/gt^-Ctrl: 1.111 ± 0.1111, and *Scn2a^gt/gt^*-hKv1.1: 1.100 ± 0.1000) (**Supplementary Figure 3I-K**). These findings indicated that hKv1.1 supplementation in the striatum and mPFC was not sufficient to affect innate behavioral deficits in adult *Scn2a*^gt/gt^ mice, possibly because these behaviors involve other brain regions or require intervention during critical developmental windows [30, 32].

## DISCUSSION

Gene therapies are advancing quickly, transforming not only how we treat diseases but also how we may cure them [33, 34]. With these advances, it is important to explore strategies to further personalize these therapies. In this study, we demonstrated doxycycline-dependent expression and functional activity of hKv1.1 in transfected ND7/23 cells and AAV-hKv1.1 injected mice. Using whole-cell patch-clamp recordings, we found that MSNs and Layer V pyramidal neurons of *Scn2a*^gt/gt^ mice exhibited intrinsic hyperexcitability. Additionally, we found that exogenous expression of hKv1.1 in *Scn2a*-deficient mice led to a reduction in the intrinsic hyperexcitability of the MSNs and Layer V pyramidal neurons. Furthermore, we observed that exogenous expression of hKv1.1 led to an increase in sociability in *Scn2a*-deficient mice; however, it did not affect innate behavioral deficits. Notably, the improvement in sociability was reversed following withdrawal of doxycycline chow, indicating that the observed improvement in social behavior was dependent on ongoing hKv1.1 expression.

In our previous study, we discovered that adult *Scn2a*^gt/gt^ mice exhibit intrinsic hyperexcitability in both striatal MSNs and Layer V pyramidal neurons of the mPFC [18]. Additionally, we demonstrated that 4TFMPG, a selective Kv1.1 opener [35], could lower neuronal excitability in striatal MSNs of *Scn2a*^gt/gt^ mice. Kv1.1 plays a crucial role in regulating neuronal excitability, as demonstrated by studies showing that application of selective Kv1.1 blockers, such as dendrotoxin-K (DTX-K), increases action potential firing and lowers rheobase [36, 37]. We expand on those findings by showing that doxycycline-induced expression of hKv1.1 was sufficient to attenuate intrinsic neuronal hyperexcitability in *Scn2a*^gt/gt^ mice. When we exogenously expressed hKv1.1 in striatal MSNs and mPFC Layer V pyramidal neurons of *Scn2a^gt/gt^* mice, we observed a decrease in the number of action potentials and an increase in rheobase. We also observed a decrease in RMP, consistent with our previous electrophysiological findings using the Kv1.1 opener 4TFMPG [18], and with other findings indicating that knocking out Kv1.1 leads to an increased RMP [19]. Overall, our findings validate the crucial role of Kv1.1 in regulating neuronal excitability.

In this study, we focused on two brain regions, the striatum and mPFC, given their involvement in different aspects of sociability, including the encoding of social sensory cues and social reward, as well as our previous finding that reinstatement of *Scn2a* in the striatum alone cannot rescue social behavior [16, 38, 39]. Our findings show that exogenous expression of hKv1.1 in *Scn2a-*deficient mice increases sociability, highlighting the importance of both of these brain regions in the manifestation of social abnormalities in *Scn2a*-related ASD. The lack of rescue of innate behavioral deficits likely reflects the selectivity of our hKv1.1 delivery to the mPFC and striatum, as behaviors such as nesting and marble burying were shown to be predominantly regulated by other regions, including the lateral hypothalamus and dorsal hippocampus [40, 41].

As gene therapies emerge as treatments for monogenic disorders, there is a strong need to optimize their application by enabling flexible titration of gene dosage. Although we did not test the minimum gene dosage required for the rescues observed, our proof-of-concept study highlights the potential of this system in *SCN2A-related* disorders. As stated previously, *SCN2A* variants generally fall into two categories: GoF and LoF. However, 20%-30% of individuals with *SCN2A*-related ASD resulting from LoF variants will develop seizures later in life [42]. In a separate study, we have shown that this virus can also mitigate absence-like seizures in a seizure-susceptible *Scn2a^gt/gt^* model [22]. These studies together suggest a potential convergent genetic intervention to treat different phenotypes in a well-defined patient population with *SCN2A* deficiency.

In conclusion, our findings demonstrate that Kv1.1 is a promising therapeutic target for *Scn2a*-related disorders and establish the feasibility of a titratable gene-therapy platform for conditions driven by neuronal hyperexcitability. Beyond its therapeutic potential, this system also provides a powerful framework for probing Kv1.1 regulation and circuit-level dynamics in genetic ASD models, laying the groundwork for future precision strategies aimed at restoring excitability balance and ameliorating behavioral deficits.

## METHODS

### Experimental animals

All experimental mice (WT (*Scn2a^+/+^*) and *Scn2a^gt/gt^*) were offspring of heterozygous (*Scn2a^+/gt^*) breeding pairs. Mice were group-housed with same-sex littermates (2-5 per cage). Mice were housed on a 12-h/12-h light/dark cycle with ad libitum access to food (2018S Teklad from Envigo) and reverse-osmosis water. Cages were housed on ventilated racks and were filled with 1/8” Bed-o-cobb bedding (Anderson, Maumee, OH, USA). A >8 g shredded puck was provided as nesting material and enrichment. Adult mice (5-11 months) of both sexes were used. All experiments were approved by the Institutional Animal Care and Use Committee (IACUC).

### Genotyping

Mice were ear-punched at the time of weaning (p21-p28), and tissue samples were collected for genotyping. DNA was extracted by dissolving samples in 50 mM NaOH for 30 minutes at 95°C. Genotyping for the tm1a trapping cassette was then performed on the extracted DNA using polymerase chain reaction (PCR) with the following primers (forward 5′ to 3′: GAGGCAAAGAATCTGTACTGTGGGG, and reverse 5′ to 3′: GACGCCTGTGAATAAAACCAAGGAA). The PCR products were 240 bp (WT) and 340 bp (gt).

### RT-qPCR

Total mRNA from the mPFC and striatum of mice injected with AAV was extracted using TRIzol reagent (Thermo Fisher Scientific, 15596018). RNA was reverse transcribed using a Maxima First Strand cDNA Synthesis Kit (Thermo Fisher Scientific, K1672). cDNA was subjected to quantitative PCR using Toyobo Thunderbird SYBR qPCR Mix (Toyobo Research Reagents, QPS-201) with the following primers: (human *KCNA1* F: ATC CGC TTG GTA AGG GTT TTT AG, R: GGC AAA GTA CAC TGC ACT AGA A. Mouse *Gapdh* F: AGT CAA GGC GAA TGG GAA G, R: AAG CAG TTG GTG GTG CAG GAT G) in a C1000 Touch PCR thermal cycler (Bio-Rad). *Gapdh* mRNA levels were used as an endogenous control for normalization using the ΔCt method [43].

### Western blot

The mPFC and striatum of mice injected with AAV-hKv1.1 were collected and flash frozen until use. Brain tissue was then sonicated in RIPA lysis buffer (Thermo Fisher Scientific, 89901) supplemented with protease and phosphatase inhibitors (Thermo Fisher Scientific, A32953). Samples were then centrifuged at 16,000 g for 20 min at 4°C. Protein concentration in the supernatant was determined by bicinchoninic acid assay. Proteins in 1 × sample buffer [62.5 mM Tris-HCl (pH 6.8), 2% (w/v) SDS, 5% glycerol, 0.05% (w/v) bromophenol blue] were denatured by boiling at 95°C for 5 min. For each sample, 40 μg of total protein was loaded onto the 12% sodium dodecyl sulfate-polyacrylamide (SDS-PAGE) gels and transferred onto a PVDF membrane (Millipore, IPFL00010) by electrophoresis. Blots were blocked in 5% nonfat milk in Tris-buffered saline and Tween 20 (TBST) for 1 h at room temperature and probed with the primary antibodies against Kv1.1 (Alomone, APC, 009, 1:200) and Vinculin (ThermoFisher Scientific, PA5-29688, 1:1000) overnight at 4°C. After overnight incubation, the blots were washed three times in TBST for 10 min, followed by incubation with secondary antibody (LI-COR, 926-32211) for 1 h at room temperature. Following three 10-min washes with TBST, the immunoreactive bands were imaged using the Odyssey® CLx Imaging System (LI-COR Biosciences) and quantitatively analyzed by densitometry with ImageJ software (NIH). Each sample was normalized to its respective Vinculin control.

### Adeno-Associated Virus (AAV) production

hKv1.1-Negative control: AAV9/NegCTRLCam with the titer of 2.75×1013 GC/mL; hKv1.1-Negative fluorescence control: AAV9/TOCitPCamTA with the titer of 1.70×1013 GC/mL; hKv1.1-Positive: AAV9/TOKv1CamTA with the titer of 1.50×1013 GC/mL are gifts from Dr. Edward Perez-Reyes (University of Virginia).

### Stereotaxic AAV injection

For intracranial injections of AAV, mice were anesthetized using ketamine/xylazine (100/10 mg/kg). Once the righting reflex and response to a painful stimulus were absent, mice were secured in a stereotaxic apparatus using ear bars (RWD Ltd, China). The skulls of the mice were exposed by making a small incision. Once the skull was exposed, stereotaxic equipment was calibrated to the bregma. Holes over the target locations (AP +1.98 mm, ML ±0.37 mm, DV −2.00 mm), (AP +1.35 mm, ML ±1.00 mm, DV −4.50 mm), and (AP +1.35 mm, ML ±1.00 mm, DV −3.30 mm) were made using a drill. Mineral oil was first suctioned into a Micro Syringe Pipette (RWD) using a UMP3 UltraMicroPump system (World Precision Instruments). The viral suspension (1×10^12^ vg/mL) was then suctioned up, and the Micro Syringe Pipette was inserted into the target location at a rate of 120 µm/min. The viral suspension was then injected at a rate of 150 nL/min (1 µL per target site). After injecting the viral suspension, the pipette was left in place for 10 minutes before being retracted. Viral expression was allowed for at least 3 weeks while the mice recovered. The animal’s body weight and health condition were closely monitored during recovery.

### Patch-clamp recordings

For electrophysiological experiments assessing AAV-hKv1.1 rescue, AAVs were injected locally into the medial prefrontal cortex (AP +1.98 mm, ML ±0.37 mm, DV −2.00 mm), and striatum (AP +1.35 mm, ML ±1.00 mm, DV −4.50 mm), (AP +1.35 mm, ML ±1.00 mm, DV −3.30 mm) relative to bregma to achieve robust local transduction. A total volume of 1 μL was delivered per injection site at approximately 150 nL/min. For the hKv1.1 treatment group, AAV9/TOKv1CamTA was mixed 1:1 with the fluorescent reporter virus AAV9/TOCitPCamTA. For the control group, AAV9/NegCTRLCam was mixed 1:1 with AAV9/TOCitPCamTA. Patch-clamp recordings were performed specifically from fluorescent neurons to enrich for cells transduced by the injected AAVs.

#### Acute slice preparations

Electrophysiology was performed in slices prepared from 8-11-month-old *Scn2a^gt/gt^* and WT littermates. Mice were deeply anesthetized with ketamine/xylazine (100/10 mg/kg, i.p., 0.1 mL per 10 g of body weight), transcardially perfused, and decapitated to dissect brains into ice-cold slicing solution containing the following (in mM): 110 choline chloride, 2.5 KCl, 1.25 NaH_2_PO_4_, 25 NaHCO_3_, 0.5 CaCl_2_, 7 MgCl_2_, 25 glucose, 1 sodium ascorbate, 3.1 sodium pyruvate (bubbled with 95% O_2_ and 5% CO_2_, pH 7.4, 305-315 mOsm). Acute coronal slices containing mPFC and/or striatum (300 μm in thickness) were cut using a vibratome (Leica VT1200S, Germany), and incubated in the same solution for 10 min at 33°C. Then, slices were transferred to normal artificial cerebrospinal fluid (aCSF) (in mM): 125 NaCl, 2.5 KCl, 2 CaCl2, 2 MgCl_2_, 25 NaHCO_3_, 1.25 NaH_2_PO_4_, 10 glucose (bubbled with 95% O_2_ and 5% CO_2_, pH 7.4, 305-315 mOsm) at 33℃ for 10-20 min and at room temperature for at least 30 min before use. Slices were visualized under IR-DIC (infrared differential interference contrast) using a BX-51WI microscope (Olympus) with an IR-2000 camera (Dage-MTI).

#### Ex vivo electrophysiological whole-cell recordings

All somatic whole-cell patch-clamp recordings were performed from identified striatal MSNs or mPFC Layer V pyramidal neurons. The selection criteria for MSNs were based on morphological characteristics, with a medium-sized cell body presenting a polygonal or diamond shape viewed with a microscope equipped with IR-DIC optics (BX-51WI, Olympus), and numerous dendritic spines and their hyperpolarized RMP (lower than −80 mV), based on the published method [44]. Layer V pyramidal cells with a prominent apical dendrite were visually identified mainly by location, shape, and pClampex online membrane test parameters. Putative pyramidal cells in Layer V were identified based on regular spiking characteristics [11, 45]. To minimize variability, recordings were made on cells with low or high HCN expression levels, corresponding to intratelencephalic (IT) or pyramidal tract (PT) neurons, respectively. The selection criterion for PT pyramidal cells followed the published approach [46]. Briefly, the selection was based on their firing properties and AP shape (i.e., cells’ intrinsic ability to generate significant membrane-potential sags induced by both hyperpolarizing and depolarizing current injection at the soma). Only the recordings from putative PT neurons were used for further analysis.

For whole-cell current-clamp recordings, the internal solution contained (in mM): 122 KMeSO4, 4 KCl, 2 MgCl2, 0.2 EGTA, 10 HEPES, 4 Na2ATP, 0.3 Tris-GTP, 14 Tris-phosphocreatine, adjusted to pH 7.25 with KOH, 295-305 mOsm. The sag ratio, input resistance, and number of action potentials were obtained in response to a series of 400-ms hyperpolarizing and depolarizing current steps from −200 pA to +400 pA in increments of 50 pA, with a sweep duration of 5 s while cells were fixed at the normal RMP or a fixed potential of −80 mV. The sag ratio was calculated with the equation:

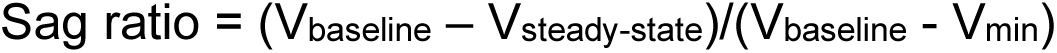

Where V_baseline_ is the resting membrane potential or −80 mV, V_min_ is the minimum voltage reached soon after the hyperpolarizing current pulse, and V_steady-state_ (Vss) is the voltage recorded at 0-10 ms before the end of the −200 pA stimulus.

The input resistance was calculated with the equation:

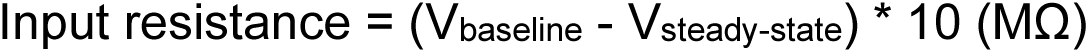

Where V_baseline_ is the resting membrane potential or −80 mV, and V_steady-state_ (V_ss_) is the voltage recorded at 0-10 ms before the end of the −100 pA stimulus.

RMP, AP threshold, amplitude, fast after-hyperpolarization (AHP), and half-width were measured from an intact AP evoked by the smallest 20 ms current step sufficient to elicit an action potential. Each sweep lasted 1.5 s, with a 10 s start-to-start interval, while cells were fixed at their normal RMP or at −80 mV. The RMP, AP threshold, amplitude, AHP, and half-width values were analyzed using the Clampfit 11.1 inbuilt statistics measurements program (criteria included the baseline, peak amplitude, antipeak amplitude, and half-width). The AP threshold was defined as the membrane potential (Vm) at which the first derivative of voltage (dV/dt) exceeded 15 V/s, and δ-AHP was calculated as the difference between the after-hyperpolarization amplitude and the baseline membrane potential.

We used thin-wall borosilicate pipettes (BF150-110-10) with open-tip resistances of 3-5 MΩ. All current-clamp recordings started at least 1 min after break-in to stabilize the contact between the glass electrode and the cell membrane, and finished within 10 min to avoid large voltage changes due to the internal solution exchange equilibrium. Recordings were performed with an Axon MultiClamp 700B amplifier (Molecular Devices), and data were acquired using pClamp 11.1 software at the normal RMP or a fixed potential of −80 mV, filtered at 2 kHz and sampling rate at 20 kHz with an Axon Digidata 1550B plus HumSilencer digitizer (Molecular Devices). Slices were maintained under continuous perfusion of aCSF at 32-33°C with a 2-3 mL/min flow. In the whole-cell configuration, recordings with stable series resistance (Rs) of 15-30 MΩ were used, and recordings with unstable Rs or a change of Rs > 20% were aborted.

### Cultured ND7/23 cells

Whole-cell patch-clamp was performed on transiently transfected ND7/23 cells (Millipore-Sigma, #92090903). Cells were transfected with Lipofectamine 2000 and 1 μg of Kv1.1 plasmid plus 0.35 μg of GFP-expressing plasmid (Green Lantern). Doxycycline was added to the medium after transfection. Potassium currents were recorded from GFP-positive cells 1-2 days after transfection.

### Open field test

On the day of the test, mice were first acclimated to the test room for 20 minutes. Mice were placed in a square arena (40 cm x 40 cm x 40 cm, Maze Engineers) for 10 min at 60 lux. Distance traveled and velocity were recorded by EthoVision XT (Noldus).

### Three-chamber assay

Social preference was assessed using a standard three-chamber choice test between an empty cylinder and a cylinder containing a same-sex novel mouse of similar age and size (40.5 cm x 60 cm x 22 cm, Maze Engineers). Mice were habituated to the three-chamber apparatus one day before the test. On the day of the test, mice were first acclimated to the test room for 20 minutes. The mice were then recorded for five minutes using EthoVision XT (Noldus). This assay was repeated under three different conditions two weeks apart from each other, while mice were maintained on control chow (Inotiv, TD.160328), 100 ppm doxycycline chow (Inotiv, TD.130840), and after returning to control chow. Videos were then hand-scored by a blinded, well-trained researcher for the number of interactions between the test mouse and the novel mouse.

### Marble burying

A clean cage was filled with 5 cm of bedding and 15 black glass marbles (1.5 cm diameter) arranged 3 by 5. Mice were placed into this cage and allowed to freely interact with the marbles for 30 minutes. The number of marbles more than two-thirds buried was recorded.

### Nesting

Mice were singly housed at the start of the dark cycle and provided a 5 cm x 5 cm nestlet (Ancare). After 36 h, the quality of the nest was analyzed. Nests were scored from 0 (nesting material untouched) to 5 (cocoon-shaped) based on nest quality and height [47].

## STATISTICAL ANALYSES

Normality, equality of variance, and outliers were assessed for all data using GraphPad Prism. For comparisons between two groups, an unpaired t-test, Welch’s t-test (parametric), or Mann–Whitney test (non-parametric) was used. For comparisons between more than two groups, one-way ANOVA with Tukey’s multiple comparisons test (parametric) or Kruskal–Wallis test with Dunn’s multiple comparisons test (non-parametric) was used. For comparisons involving two factors, two-way ANOVA with Tukey’s or Bonferroni multiple comparisons test was used.

Post hoc comparisons were carried out only when the primary measure showed statistical significance. All data were expressed as mean ± SEM, with statistical significance determined at p-values < 0.05. In all figures, p ≥ 0.05 is indicated as not significant (ns), p < 0.05 is indicated as one asterisk (*), p < 0.01 is indicated as two asterisks (**), p < 0.001 is indicated as three asterisks (***), and p < 0.0001 is indicated as four asterisks (****).

## ACKNOWLEDGMENTS

The research reported in this publication was supported by the NINDS of the NIH (R56NS097726 to EPR, R01NS117585 and R01NS123154 to Y.Y.). The authors gratefully acknowledge support from the Familie*SCN2A* Foundation and PIIN for additional funding support. Behavioral assays were performed using the Purdue Animal Behavior Core.

## AUTHOR CONTRIBUTIONS

B.D., J.Z., and Y.Y. designed research; B.D., J.Z., I.V., R.G., and M.H. performed experiments; B.D., J.Z., I.V., R.G., M.H., P.M., K.L., A.A., K.F., and D.R. analyzed data; A.A. and R.R. assisted with project administration; B.D., J.Z., Z.Z., and Y.Y. wrote the manuscript with input from E.P.R.

## COMPETING INTERESTS

The authors report no financial interests or potential conflicts of interest.

## RESOURCE AVAILABILITY

Further information and requests for resources and reagents should be directed to and will be fulfilled by the lead contact, Yang Yang.

## Figure Legends

**Figure S1:**
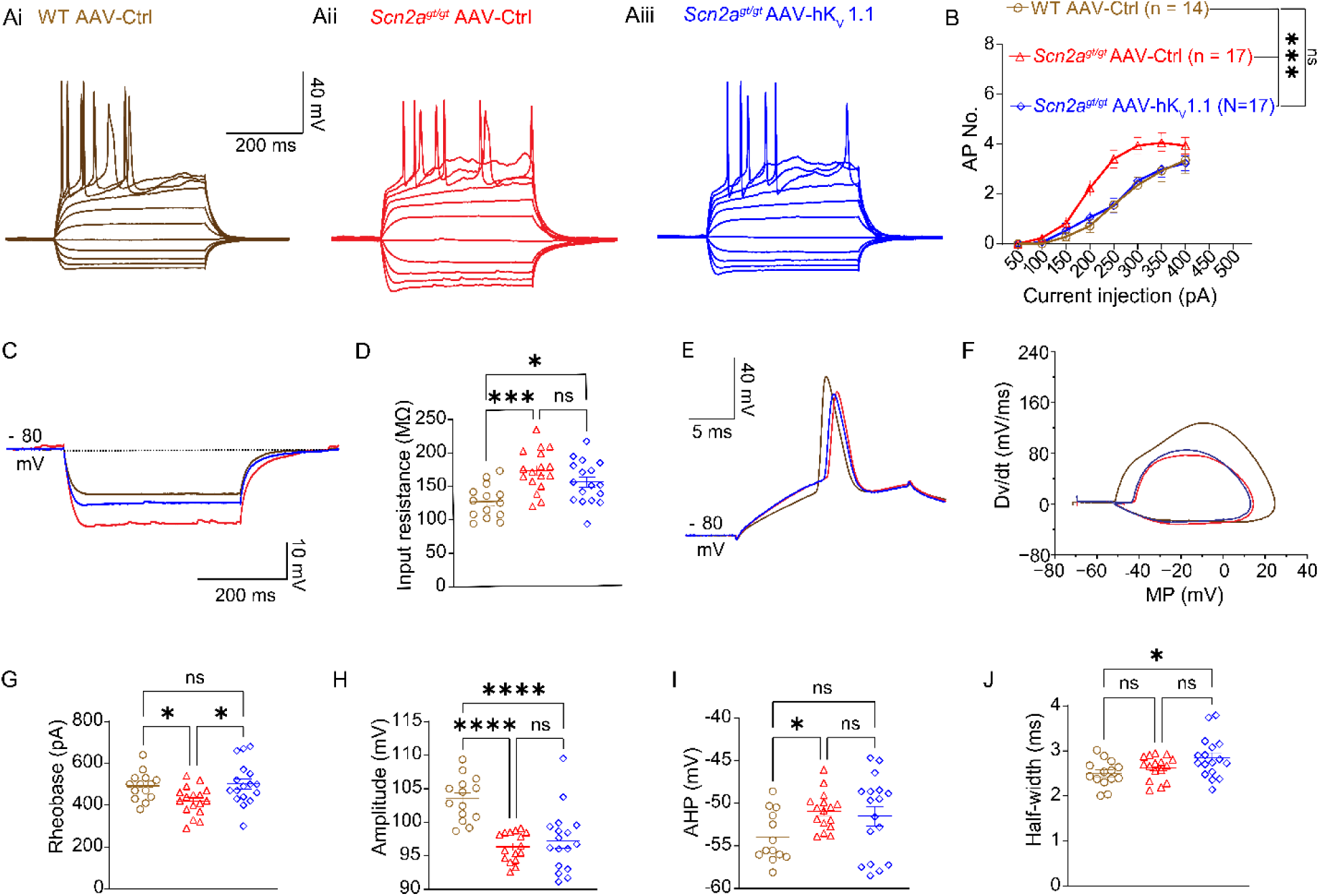
Exogenous expression of hKv1.1 in the striatum decreases neuronal firing of medium spiny neurons fixed at −80 mV in *Scn2a*-deficient mice. (Ai-Aiii) Representative current-clamp recordings were obtained from MSNs at a fixed membrane potential of −80 mV in the nucleus accumbens of WT mice injected with AAV-Ctrl (brown) and *Scn2a*-deficient mice injected with AAV-Ctrl (red) or AAV-hKv1.1 (blue). (B) Number of action potentials generated in response to depolarizing current pulses (50-400 pA) when MSNs of WT mice injected with AAV-Ctrl (brown) and *Scn2a*-deficient mice injected with AAV-Ctrl (red) or AAV-hKv1.1 (blue) are fixed at −80 mV. Two-way ANOVA with Tukey’s multiple comparisons: ns= not significant, p>0.05; ***, p<0.001. (C) Representative traces of MSNs fixed at −80 mV of WT mice injected with AAV-Ctrl (brown) and *Scn2a*-deficient mice injected with AAV-Ctrl (red) or AAV-hKv1.1 (blue) generated in response to a −100 pA current injection. (D) Input resistance of MSNs fixed at −80 mV of WT mice injected with AAV-Ctrl (brown) and *Scn2a*-deficient mice injected with AAV-Ctrl (red) or AAV-hKv1.1 (blue). One-way ANOVA with Tukey’s multiple comparisons: ns= not significant, p>0.05; *, p<0.05; ***, p<0.001. (E) Representative action potential spike from MSNs fixed at −80 mV of WT mice injected with AAV-Ctrl (brown) and *Scn2a*-deficient mice injected with AAV-Ctrl (red) or AAV-hKv1.1 (blue). (F) Action potential phase plots from MSNs of WT mice injected with AAV-Ctrl (brown) and *Scn2a*-deficient mice injected with AAV-Ctrl (red) or AAV-hKv1.1 (blue). (G) Rheobase of MSNs fixed at −80 mV of WT mice injected with AAV-Ctrl (brown) and *Scn2a*-deficient mice injected with AAV-Ctrl (red) or AAV-hKv1.1 (blue). Kruskal-Wallis with Dunn’s multiple comparisons: ns= not significant, p>0.05; *, p<0.05. (H) Action potential amplitude of MSNs of WT mice injected with AAV-Ctrl (brown) and *Scn2a*-deficient mice injected with AAV-Ctrl (red) or AAV-hKv1.1 (blue). One-way ANOVA with Tukey’s multiple comparisons: ns= not significant, p>0.05; ****, p<0.0001. (I) Fast after-hyperpolarization (AHP) of action potential spikes of MSNs from WT mice injected with AAV-Ctrl (brown) and *Scn2a*-deficient mice injected with AAV-Ctrl (red) or AAV-hKv1.1 (blue). One-way ANOVA with Tukey’s multiple comparisons: ns= not significant. (J) Action potential half-width of MSNs from WT mice injected with AAV-Ctrl (brown) and *Scn2a*-deficient mice injected with AAV-Ctrl (red) or AAV-hKv1.1 (blue). One-way ANOVA with Tukey’s multiple comparisons: ns= not significant.

**Figure S2:**
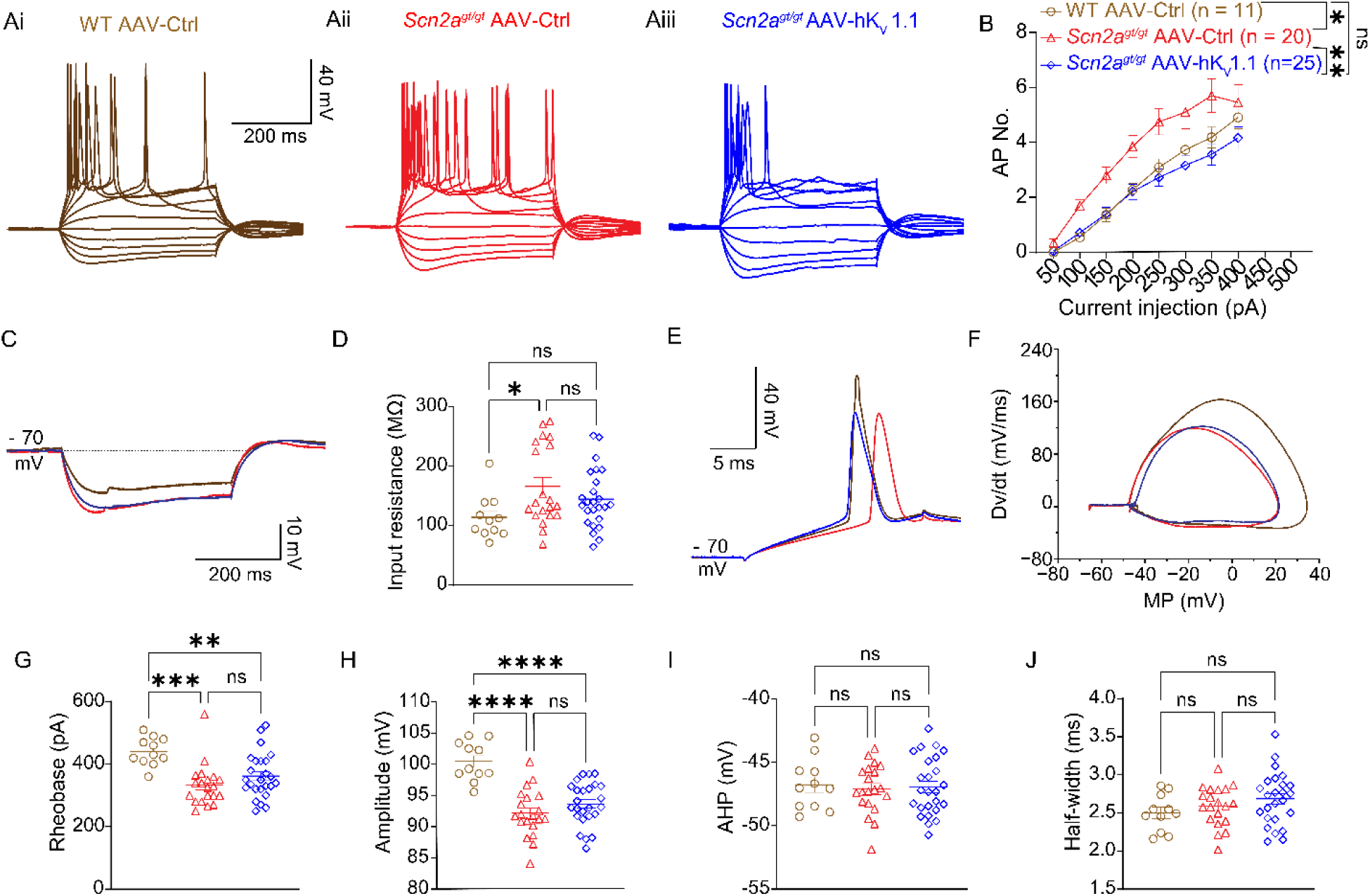
Exogenous expression of hKv1.1 in the medial prefrontal cortex decreases neuronal firing of Layer V pyramidal neurons fixed at −70 mV in *Scn2a*-deficient mice. (Ai-Aiii) Representative current-clamp recordings of Layer V pyramidal neurons obtained at a fixed membrane potential of −70 mV from WT mice injected with AAV-Ctrl (brown) and *Scn2a*-deficient mice injected with AAV-Ctrl (red) or AAV-hKv1.1 (blue). (B) Number of action potentials generated in response to depolarizing current pulses (50-400 pA) when Layer V pyramidal neurons from WT mice injected with AAV-Ctrl (brown) and *Scn2a*-deficient mice injected with AAV-Ctrl (red) or AAV-hKv1.1 (blue) are fixed at −70 mV. Two-way ANOVA with Tukey’s multiple comparisons: ns= not significant, p>0.05; *, p<0.05; **, p<0.01. (C) Representative traces of Layer V pyramidal neurons fixed at −70 mV from WT mice injected with AAV-Ctrl (brown) and *Scn2a*-deficient mice injected with AAV-Ctrl (red) or AAV-hKv1.1 (blue) generated in response to a −100 pA injection. (D) Input resistance of Layer V pyramidal neurons fixed at −70 mV from WT mice injected with AAV-Ctrl (brown) and *Scn2a*-deficient mice injected with AAV-Ctrl (red) or AAV-hKv1.1 (blue). One-way ANOVA with Tukey’s multiple comparisons: ns= not significant, p>0.05; *, p<0.05. (E) Representative action potential spike from Layer V pyramidal neurons fixed at −70 mV of WT mice injected with AAV-Ctrl (brown) and *Scn2a*-deficient mice injected with AAV-Ctrl (red) or AAV-hKv1.1 (blue). (F) Action potential phase plots from Layer V pyramidal neurons of WT mice injected with AAV-Ctrl (brown) and *Scn2a*-deficient mice injected with AAV-Ctrl (red) or AAV-hKv1.1 (blue). (G) Rheobase of Layer V pyramidal neurons fixed at −70 mV of WT mice injected with AAV-Ctrl (brown) and *Scn2a*-deficient mice injected with AAV-Ctrl (red) or AAV-hKv1.1 (blue). Kruskal-Wallis with Dunn’s multiple comparisons: ns= not significant, p>0.05; ***, p<0.001. (H) Action potential amplitude of Layer V pyramidal neurons fixed at −70 mV of WT mice injected with AAV-Ctrl (brown) and *Scn2a*-deficient mice injected with AAV-Ctrl (red) or AAV-hKv1.1 (blue). One-way ANOVA with Tukey’s multiple comparisons: ns= not significant, p>0.05; ****, p<0.0001. (I) Fast after-hyperpolarization (AHP) of action potential spikes of Layer V pyramidal neurons fixed at −70 mV from WT mice injected with AAV-Ctrl (brown) and *Scn2a*-deficient mice injected with AAV-Ctrl (red) or AAV-hKv1.1 (blue). One-way ANOVA with Tukey’s multiple comparisons: ns= not significant. (J) Action potential half-width of Layer V pyramidal neurons fixed at −70 mV from WT mice injected with AAV-Ctrl (brown) and *Scn2a*-deficient mice injected with AAV-Ctrl (red) or AAV-hKv1.1 (blue). One-way ANOVA with Tukey’s multiple comparisons: ns= not significant.

**Figure S3:**
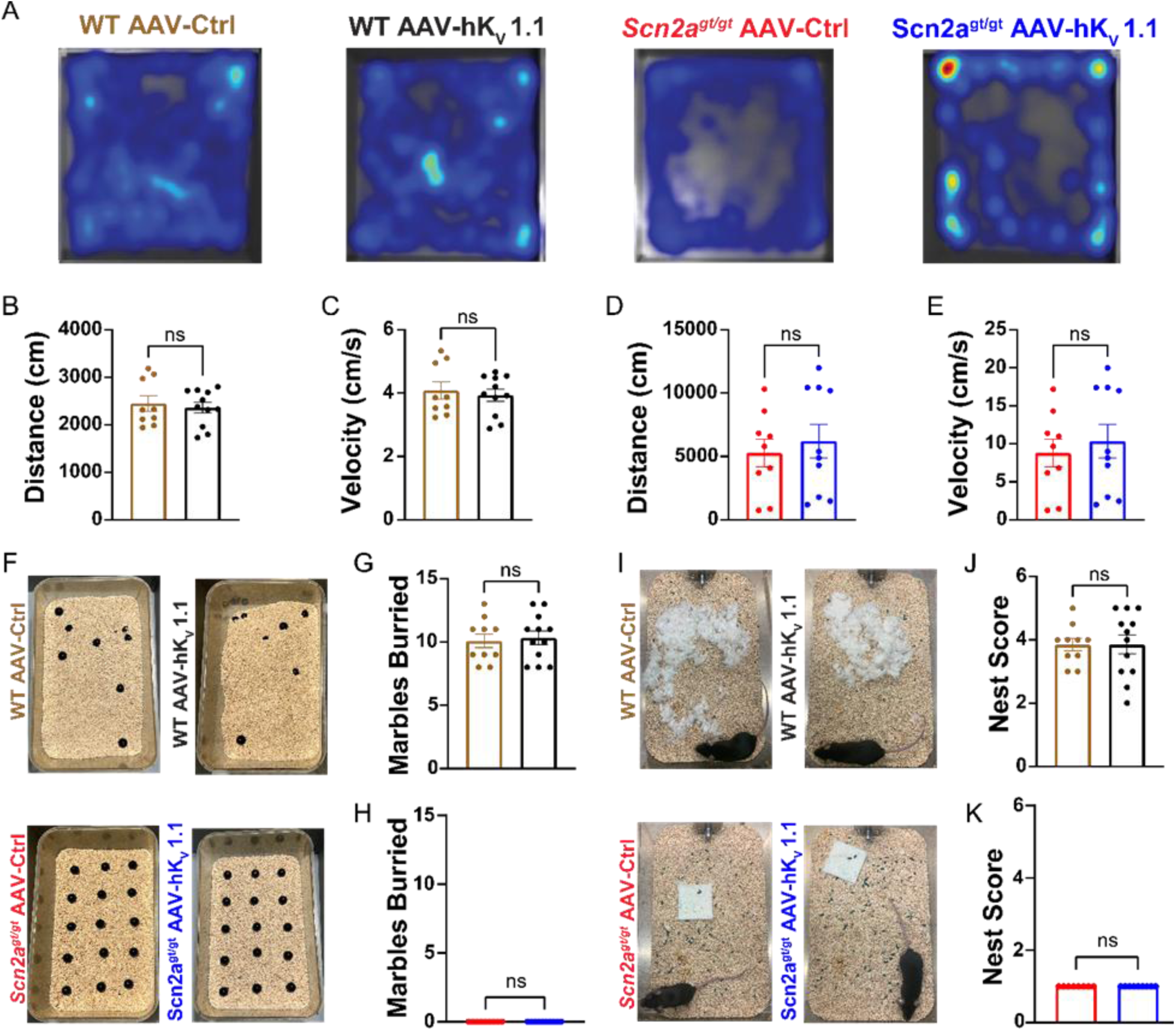
Exogenous expression of hKv1.1 does not rescue innate behavior in *Scn2a*-deficient mice. (A) Representative heatmaps of WT or *Scn2a*-deficient mice injected with AAV-Ctrl or AAV-hKv1.1 during a 10-minute open field at 60 lux. (B) Distance traveled by WT mice injected with AAV-Ctrl (brown) or AAV-hKv1.1 (black) during an open field assay. n= WT-Ctrl: 9, WT-hKv1.1: 11. Unpaired t-test: ns= not significant, p>0.05. (C) Velocity of WT mice injected with AAV-Ctrl (brown) or AAV-hKv1.1 (black) during an open field assay. n= WT-Ctrl: 9, WT-hKv1.1: 11. Unpaired t-test: ns= not significant, p>0.05. (D) Distance traveled by *Scn2a*-deficient mice injected with AAV-Ctrl (red) or AAV-hKv1.1 (blue) during an open field assay. n= *Scn2a*-Ctrl: 9, *Scn2a*-hKv1.1: 10. Unpaired t-test: ns= not significant, p>0.05. (E) Velocity of *Scn2a*-deficient mice injected with AAV-Ctrl (red) or AAV-hKv1.1 (blue) during an open field assay. n= *Scn2a*-Ctrl: 9, *Scn2a*-hKv1.1: 10. Unpaired t-test: ns= not significant, p>0.05. (F) Representative images of marbles buried after 30 minutes during the marble burying assay. (G) Marbles buried by WT mice injected with AAV-Ctrl (brown) or AAV-hKv1.1 (black) during a 30-minute marble burying assay. n= WT-Ctrl: 10, WT-hKv1.1: 12. Unpaired t-test: ns= not significant, p>0.05. (H) Marbles buried by *Scn2a*-deficient mice injected with AAV-Ctrl (red) or AAV-hKv1.1 (blue) during a 30-minute marble burying assay. n= *Scn2a*-Ctrl: 9, *Scn2a*-hKv1.1: 10. Mann-Whitney test: ns= not significant, p>0.05. (I) Representative images of nests constructed over 36 h by WT mice injected with AAV-Ctrl (brown), Scn2a-deficient mice injected with AAV-Ctrl (red), WT mice injected with AAV-hKv1.1 (black), and Scn2a-deficient mice injected with AAV-hKv1.1 (blue). (J) Nest score of WT mice injected with AAV-Ctrl (brown) or AAV-hKv1.1 (black) after 36 h. n= WT-Ctrl: 10, WT-hKv1.1: 12. Unpaired t-test: ns= not significant, p>0.05. (K) Nest score of *Scn2a*-deficient mice injected with AAV-Ctrl (red) or AAV-hKv1.1 (blue) after 36 h. n= *Scn2a*-Ctrl: 9, *Scn2a*-hKv1.1: 10. Unpaired t-test: ns= not significant, p>0.05.

## Notes

### Competing Interest Statement

The authors have declared no competing interest.

